# Novelty-related engagement of VTA and anterior hippocampus propagate changes in cortical network plasticity at different scales

**DOI:** 10.1101/2021.03.22.436448

**Authors:** Emily T. Cowan, Matthew Fain, Ian O’Shea, Lauren M. Ellman, Vishnu P. Murty

**Affiliations:** Temple University, Philadelphia, PA, 19122; University of California, San Diego, La Jolla, CA 92093

## Abstract

The detection of novelty indicates changes in the environment and the need to update existing representations. In response to novelty, interactions across the ventral tegmental area (VTA)-hippocampal circuit support experience-dependent plasticity in the hippocampus. While theories have broadly suggested plasticity-related changes are also instantiated in the cortex, research has also shown evidence for functional heterogeneity in cortical networks. It therefore remains unclear how the hippocampal-VTA circuit engages cortical networks, and whether novelty targets specific cortical regions or diffuse, large-scale cortical networks. To adjudicate the role of the VTA and hippocampus in cortical network plasticity, we used human functional magnetic resonance imaging (fMRI) to compare resting state functional coupling before and following exposure to novel scene images. Functional coupling between right anterior hippocampus and VTA was enhanced following novelty exposure. However, we also found evidence for a double dissociation, with anterior hippocampus and VTA showing distinct patterns of post-novelty functional coupling enhancements, targeting task-relevant regions versus large-scale networks, respectively. Further, significant correlations between these networks and the novelty-related plasticity in the anterior hippocampal-VTA functional network suggest the central hippocampal-VTA network may facilitate the interactions with the cortex. These findings support an extended model of novelty-induced plasticity, in which novelty elicits plasticity-related changes in both local and global cortical networks.

**Significance Statement:** Novelty detection is critical for adaptive behavior, signaling the need to update existing representations. By engaging the bi-directional hippocampal-VTA circuit, novelty has been shown to induce plasticity-related changes in the hippocampus. However, it remains an open question how novelty targets such plasticity-related changes in cortical networks. We show that anterior hippocampus and VTA target cortical networks at different spatial scales, with respective enhancements in post-novelty functional coupling with a task-relevant cortical region and a large-scale memory network. The results presented here support an extended model of novelty-related plasticity, in which engaging the anterior hippocampal-VTA circuit through novelty exposure propagates cortical plasticity through hippocampal and VTA functional pathways at distinct scales, targeting specific or diffuse cortical networks.

## Introduction

Novelty indicates that existing representations of the world need to be updated with new information. The brain has specialized systems to process novelty, resulting in plasticity-related changes that facilitate the persistence of new information (Ranganath and Rainer, 2003; Lisman and Grace, 2005; Shohamy and Adcock, 2010). Prior work has focused on how circuits centered on the hippocampus and VTA contribute to novelty-related plasticity, yet, open questions remain as to how novelty restructures functional networks with the cortex.

The VTA-hippocampal circuit seems to be critical in novelty processing. The hippocampus has been shown to detect novelty (Knight, 1996; Tulving et al., 1996; Strange et al., 1999; Ranganath and Rainer, 2003; Axmacher et al., 2010; Shohamy and Adcock, 2010; Kafkas and Montaldi, 2018), and in response, signals to the VTA to stimulate the release of dopamine (Lisman and Grace, 2005), which enhances hippocampal synaptic plasticity (Huang and Kandel, 1995; Li et al., 2003; Lisman and Grace, 2005; Moncada and Viola, 2007; Rossato et al., 2009; Bethus et al., 2010; Shohamy and Adcock, 2010). Hippocampal-VTA interactions are therefore critical in the selective strengthening of synapses representing novel information, facilitating its retention. However, less attention has focused on how engagement of this circuit promotes experience-dependent cortical plasticity in humans.

The neocortex plays a critical role in the persistent representation of memories. Theories suggest that, while the hippocampus rapidly encodes new information, the cortex has a slower learning rate, gradually extracting the central information from the hippocampus through the repeated reactivation of the trace (McClelland et al., 1995), a phenomenon known as replay (Wilson and McNaughton, 1994; Girardeau and Zugaro, 2011; Joo and Frank, 2018). The newly acquired information therefore becomes more abstractly represented and can be incorporated into existing cortical stores without interference (McClelland et al., 1995; Moscovitch et al., 2016). Dopamine influences replay, such that optogenetic stimulation of dopaminergic VTA-hippocampal projections during novelty exposure enhances subsequent replay (McNamara et al., 2014). It follows that through hippocampal-cortical interactions existing cortical representations can also be updated with novel information (Squire and Zola-Morgan, 1991; Wang and Morris, 2010; Moscovitch et al., 2016). However, research has yet to consider which cortical networks VTA-hippocampal circuits target for plasticity-related changes.

Theories predicated on hippocampal-cortical interactions tend to treat cortex as one homogenous structure. In contrast, research has shown that replay is coordinated between the hippocampus and specific neocortical regions (Ji and Wilson, 2007; Lansink et al., 2009; Peyrache et al., 2009; Wierzynski et al., 2009) and provided evidence for functional differences in cortical networks (Ritchey et al., 2015; Barnett et al., 2020). It remains unclear how effects of novelty would map onto such functional heterogeneity in cortical network organization. One possibility is that novelty facilitates plasticity-related changes in networks with relatively specific cortical targets, affecting only those regions specialized for processing the content of the novel information. Alternatively, novelty-related plasticity could target a widespread network of cortical regions, eliciting diffuse effects on cognitive processes. Evidence for both possibilities exist, albeit through different pathways. Reports have shown experience-dependent enhancements in functional coupling specifically between hippocampus and task-relevant cortical regions (Tambini et al., 2010; Vilberg and Davachi, 2013; Schlichting and Preston, 2014; Murty et al., 2017b; Collins and Dickerson, 2019), whereas VTA dopaminergic signaling has widespread projections across regions beyond the hippocampus (Shohamy and Adcock, 2010; Murty and Dickerson, 2016).

The aim of the current study was to interrogate how the hippocampus and VTA restructure network dynamics for plasticity-related changes following novelty exposure. Using fMRI, we compared functional coupling during resting state scans prior to, and following, exposure to novel scene images. We report evidence for a double dissociation in the scale of novelty-related functional coupling enhancements, with right anterior hippocampus and VTA respectively only showing effects with a task-relevant cortical region and large-scale memory network. These findings support a model by which engaging the hippocampal-VTA circuit through novelty exposure propagates changes in cortical networks at different scales of distribution.

## Materials and Methods

### Participants

Participants were recruited for this experiment as control subjects in a larger study examining psychosis risk. The final sample with usable data on the novelty task and pre- and post-task resting state scans yielded thirty-seven participants included in all analyses. Informed consent was obtained from each participant in a manner approved by Temple University’s Institutional Review Board.

### Procedures

The protocol and materials utilized here were based on previously published work (Murty et al., 2013, 2017a). In brief, the task involved two phases, a familiarization and a novelty exposure phase (Figure 1). Participants first completed the familiarization phase in which outdoor scene images were shown one at a time while participants completed a continuous recognition task; 80 scene images were shown six times (“familiar”) and 40 were shown once (foils). Approximately 20 minutes later participants entered the MRI scanner for the novelty exposure phase, in which they again viewed a sequence of outdoor scene images while completing a target detection task, pressing a button when a specific target scene image was shown. The presented images included 80 “novel” images that had never been seen before, 80 “familiar” images (those previously viewed six times during the familiarization phase), and the “target” scene image (40 repeated presentations), in a randomized order.

**Figure 1.**
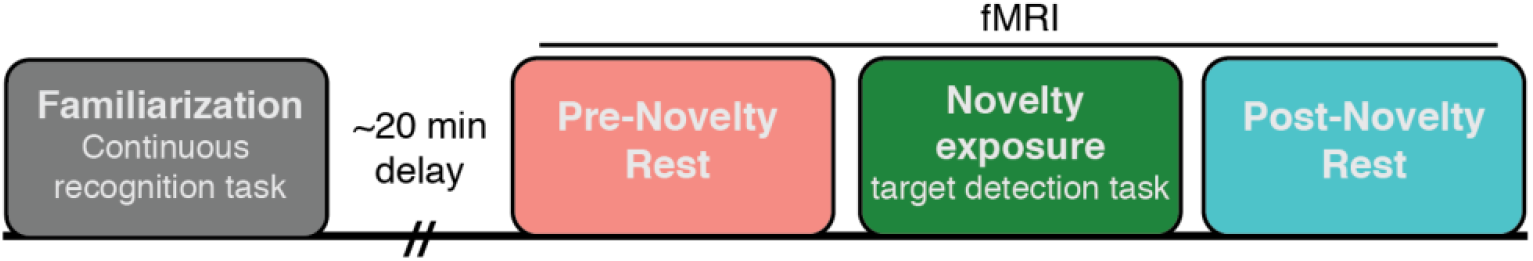
Experimental Design. In the first phase of the experiment, participants were familiarized with scene images during a continuous recognition task. In the scanner, participants first underwent a baseline resting state scan (pre-novelty rest), followed by the novelty-exposure phase, in which novel scene images were presented intermixed with the familiar images while participants completed a target detection task. Immediately following this task, participants completed a second, post-novelty, resting state scan.

Critically, and where the current experiment diverges from prior published work using this protocol, all participants completed resting state scans prior to and following the target detection task in the fMRI scanner, a design used previously to compare experience-dependent changes in resting state functional coupling (Tambini et al., 2010; Tompary et al., 2015; Murty et al., 2017b). Resting state sessions lasted 5.8 minutes, and participants were instructed to keep their eyes open and look at the fixation cross on the screen.

### MRI data acquisition and preprocessing

Scanning was completed on a 3T Siemens Magnetom Prisma scanner. Functional imaging data was collected using a multi-band echo-planar (EPI) pulse sequence (TR= 1.73s, TE= 25, voxel size= 2.38 × 2.38 × 2.5mm, MB factor: 2). In addition, a high-resolution T1 weighted anatomical scan (magnetization-prepared rapid-acquisition gradient echo sequence, voxel size= 0.9mm isotropic) was acquired to aid in functional image co-registration.

All fMRI preprocessing was performed using the FSL (version 6.0.3) fMRI Expert Analysis Tool version 6 (FSL: http://fsl.fmrib.ox.ac.uk/fsl/). Functional images were skull stripped with the Brain Extraction Tool, high-pass filtered (60s cutoff), spatially smoothed with a 5mm FWHM kernel, intensity normalized, and MCFLIRT was applied for motion correction. Functional data was registered to the high-resolution anatomical scans with FSL’s FLIRT tool (Linear with BBR), then to standard Montreal Neurological Institute (MNI) space using FNIRT’s nonlinear registration (12 DOF, 10mm warp resolution). Additionally, noise-related measures were computed for average signal in CSF and white matter masks (generated using FSL’s FAST segmentation tool), time points of excessive head motion (identified using FSL’s motion outliers tool), as well as the six head motion parameters and their first derivatives. The same preprocessing pipeline was carried out for the rest and task scans.

### Region of Interest (ROI) definition

Anatomical hippocampal ROIs were defined using the Harvard Oxford Subcortical Probabilistic Atlas, thresholded at 50%, and in MATLAB, divided lengthwise into anterior, mid, and posterior thirds using custom code. The VTA ROI was defined based on a probabilistic atlas, thresholded at 75% (Murty et al., 2014). Additional ROIs were defined based on coordinates reported in prior work using the exact same task; for the parahippocampal cortex ROI, we used the coordinates for a region in left PhC previously shown to be sensitive to novelty exposure (Murty et al., 2013) (MNI coordinates: −28, −54, −14). The ROIs in the PMAT network were based on coordinates reported in a previous publication defining these two networks (Ritchey et al., 2014), though we excluded the hippocampal ROIs included in the PMAT network as we were interested in dynamics between hippocampus and VTA with these regions. The full list of PMAT ROIs is included in Table 1. All coordinates were used to generate a spherical ROI using a 5mm kernel.

**Table 1.** Coordinates for PMAT ROIs. The coordinates for the PMAT ROIs were adapted from Ritchey et al., 2014, and used as central points to generate 5mm spherical kernels. Shaded rows represent the ROIs in the AT network, while non-shaded rows make up the PM network.

|  | **Region** | **MNI coordinates**<br>**(x, y, z)** |
| --- | --- | --- |
| 1 | RPTHAL4 | 22, -30, 6 |
| 2 | LMOCC1 | -2, -78, -2 |
| 3 | LOCCP1 | -16, -96, 22 |
| 4 | ROCC2 | 16, -98, 20 |
| 5 | RPREC3 | 18, -68, 24 |
| 6 | RMOCC3 | 14, -72, 8 |
| 7 | ROCCP2 | 14, -88, 4 |
| 8 | LPHC1 | -14, -50, -6 |
| 9 | RPHC2 | 18, -46, -4 |
| 10 | LPREC1 | -14, -60, 18 |
| 11 | RMOCC2 | 6, -58, 14 |
| 12 | LOCC1 | -38, -82, 28 |
| 13 | RANG2 | 52, -48, 28 |
| 14 | LPREC5 | -2, -60, 34 |
| 15 | LRSC1 | -8, -50, 14 |
| 16 | LPREC2 | -12, -50, 40 |
| 17 | RRSC2 | 8, -46, 16 |
| 18 | RPREC4 | 18, -52, 36 |
| 19 | LDLPFC1 | -24, 60, 24 |
| 20 | LMPFC | -2, 60, 34 |
| 21 | RDLPFC2 | 18, 58, 24 |
| 22 | RTPC1 | -44, 4, -42 |
| 23 | RPMTG2 | 54, -2, -32 |
| 24 | ROFC2 | -6, 16, -22 |
| 25 | ROFC3 | 24, 12, -24 |
| 26 | RTPC2 | 38, 20, -40 |
| 27 | LOFC1 | -16, 24, -20 |
| 38 | LPMTG1 | -64, -36, -10 |
| 29 | RAITG2 | 64, -14, -28 |
| 30 | RFPC2 | 38, 60, -10 |
| 31 | RFUS2 | 40, -18, -28 |
| 32 | RPRC | 30, -12, -36 |
| 33 | LFUS1 | -42, -14, -30 |
| 34 | ROFC4 | 8, 22, -20 |

### fMRI data analysis

#### Functional coupling analysis

To compute resting state functional coupling measures, we first calculated GLMs for the pre- and post-novelty exposure resting state runs, which only included noise regressors. The data were then registered to standard MNI space and bandpass filtered between 0.01 and 0.1. For both runs we then extracted the average timecourse from each ROI. We measured functional coupling as the correlation between the timecourses between ROIs. Functional coupling between pairs of ROIs was calculated as the correlation between each ROI’s timecourse (e.g., for anterior hippocampus and VTA, or anterior hippocampus with PhC), while the functional coupling with the PMAT network ROIs was calculated by taking the average of each pairwise correlation between the seed region (anterior hippocampus or VTA) with each ROI in the PMAT network. We calculated both the coupling with the combined PMAT network, as well as the coupling with the respective PM and AT regions. Fisher’s r-to-z transformations were applied to all pairwise correlation measures before averaging and further statistical analyses were computed.

#### Univariate analyses during novelty-exposure task

To measure BOLD response during the novelty exposure phase of the task, we computed a GLM with three regressors for the trials of the novel, familiar, and target conditions. Following on our prior work, this GLM also included three regressors measuring the habituation for each trial type, calculated as a linear decrease across the trial presentations. Noise-related measures (see above) were also added as additional nuisance regressors. The resulting contrasts were registered to standard MNI space, from which we then extracted the beta parameters from each contrast of interest (novel and familiar greater than baseline, novel > familiar). We focused these analyses on the anterior and posterior hippocampus and VTA ROIs, as our central question with these analyses was to examine whether novelty-evoked univariate activation correlated with the change in functional coupling from pre to post-novelty exposure.

#### Statistical analysis

All reported statistical analyses are two-tailed, and *P* < 0.05 was considered significant for all tests. Repeated measures ANOVAs were performed and followed up with paired sample *t*-tests where applicable. Statistics were performed with R version 3.5.1, RStudio (RStudio, Inc version 0.99.903) and MATLAB (MathWorks) using both built-in and custom functions. Bonferroni correction was applied to the four tests used to assess change in hippocampal-VTA functional coupling calculated for long-axis region and hemisphere using p.adjust in R.

## Results

### Hippocampal-VTA functional connectivity

The circuit between the hippocampus and VTA is critical in novelty processing, from the detection of novelty to the subsequent dopaminergic-based enhancements in LTP in the hippocampus (Lisman and Grace, 2005; Shohamy and Adcock, 2010). As such, we predicted that functional coupling between the hippocampus and VTA would be enhanced following the novelty exposure session. We expected to see this relationship particularly between VTA and anterior, rather than posterior, hippocampus as the former has more often been implicated in novelty processing (Strange et al., 1999; Poppenk et al., 2013; Kafkas and Montaldi, 2018). We additionally particularly focused on right hippocampus, as prior work using this paradigm found a peak in novelty-related activation in this region (Murty et al., 2013). Pre- and post-task measures of functional coupling between anterior and posterior hippocampus with VTA were separately calculated (see Methods).

As seen in Figure 2a, the functional coupling between right anterior hippocampus and VTA was significantly greater during the post-task compared to pre-task session (*t*(36) = −3.07, p= 0.004; Bonferroni corrected p= 0.016). However, as predicted, right posterior hippocampal-VTA functional coupling did not significantly differ across resting state scans (*t*(36) = −1.34, p = 0.19, corrected p= 0.75). Left anterior and posterior hippocampal functional coupling with VTA did not show significant changes across sessions (left anterior, *t*(36)= −1.69, p= 0.1, corrected p= 0.39; left posterior *t*(36)= −1.66, p= 0.11, corrected p= 0.42). Together, these results illustrate that functional coupling between anterior hippocampus and VTA is sensitive to novelty-related plasticity enhancements.

**Figure 2.**
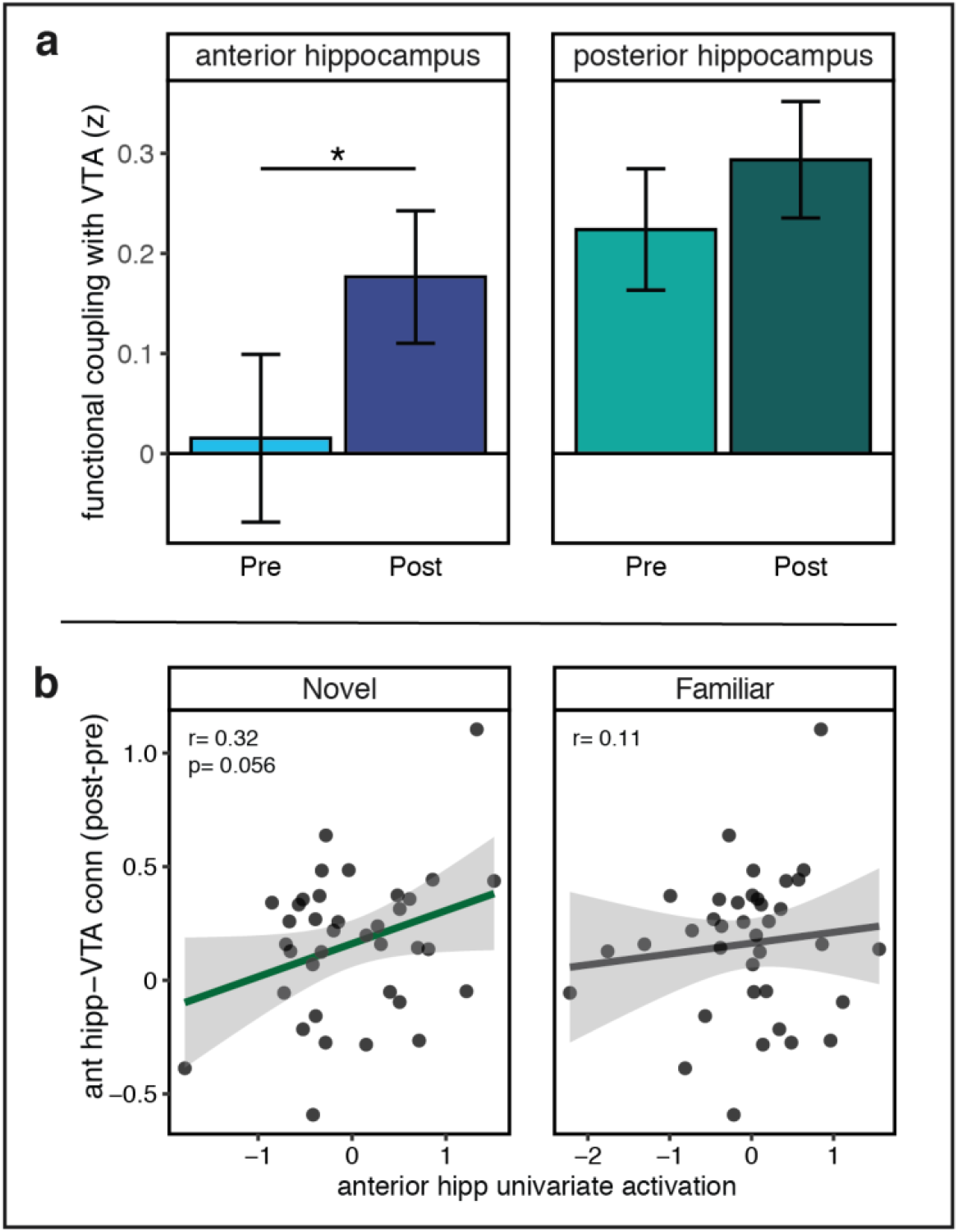
Resting state functional coupling between hippocampus and VTA. a) Right anterior hippocampal-VTA functional coupling is significantly greater during the post-novelty exposure rest scan (dark blue) compared to the pre-novelty baseline (light blue), while posterior hippocampal-VTA functional coupling did not significantly change following novelty exposure. b) The change in right anterior hippocampal-VTA functional coupling (post – pre-novelty exposure) marginally correlates with univariate activation in right anterior hippocampus during novel, but not familiar, trials of the novelty-exposure phase.

The hippocampal-VTA loop is thought to rely on the hippocampus’ role as a novelty detector. It is therefore expected that the enhancement in anterior hippocampal-VTA functional coupling would be related to the responsiveness of the anterior hippocampus to novelty, measured as univariate activation during novel trials in the novelty-exposure task (see Methods). The change in right anterior hippocampal-VTA functional coupling, calculated as the difference between functional coupling during the post- and pre-novelty exposure rest sessions, marginally correlated with the univariate activation in right anterior hippocampus during novel trials (*r*= 0.32, p= 0.056; Figure 2b). This relationship with univariate activation in anterior hippocampus during novel trials is consistent with prior work suggesting the circuit between the hippocampus and VTA is activated via the hippocampus’ role as a novelty detector (Lisman and Grace, 2005). There was not a significant relationship between hippocampal activation during familiar trials and changes in VTA-anterior hippocampal coupling (*r*= 0.11, p= 0.51).

Together, these results confirm the importance of interactions between the hippocampus and VTA in novelty processing, providing evidence that plasticity-related changes in the hippocampal-VTA circuit are related to the response to the exposure to novelty in the hippocampus.

### Dissociation in hippocampal and VTA plasticity-related targets

The main goal of this experiment was to query the scale by which novelty-related plasticity changes restructure cortical network dynamics; targeting local task-relevant sensory cortical regions versus global, memory-related network of regions. We therefore examined changes in functional coupling between right anterior hippocampus and VTA with the task-relevant parahippocampal cortex (PhC), a region known to process scene images and scene novelty (Davachi, 2006; Howard et al., 2011; Murty et al., 2013), and the more widespread PMAT network, which consists of both the posterior medial (PM) and anterior temporal (AT) networks known to be involved in memory processing (Ranganath and Ritchey, 2012; Ritchey et al., 2015) (see Methods). A 2(Seed: ant hipp, VTA) × 2(Network: PMAT, PhC) × 2(Session: Pre, Post) repeated measures ANOVA yielded a significant Seed × Network × Session interaction effect (*F*(1,36)= 9.16, p=0.005), indicating a dissociation in the experience-dependent changes in functional coupling between anterior hippocampus and VTA depending on the target cortical network. This analysis also resulted in a significant Seed × Network interaction (*F*(1,36)=13.2, p=0.0009), and a marginal main effect of session (*F*(1,36)=3.8, p=0.059).

To unpack the significant interaction, we conducted follow up 2(Seed: aHPC, VTA) × 2(Session: Pre, Post) repeated measures ANOVAs for each network target, PhC and PMAT, separately. For PhC, the ANOVA resulted in a main effect of Seed region (*F*(1,36)=5.55, p= 0.024), and a significant Seed × Session interaction (*F*(1,36)= 6.21, p=0.017). As shown in Figure 3a, follow up t-tests revealed that this significant interaction effect was driven by a post-encoding enhancement in anterior hippocampal-PhC functional coupling (*t*(36)= −2.67, p= 0.01), with no significant change in VTA-PhC functional coupling (*t*(36)=0.28, p= 0.78).

**Figure 3.**
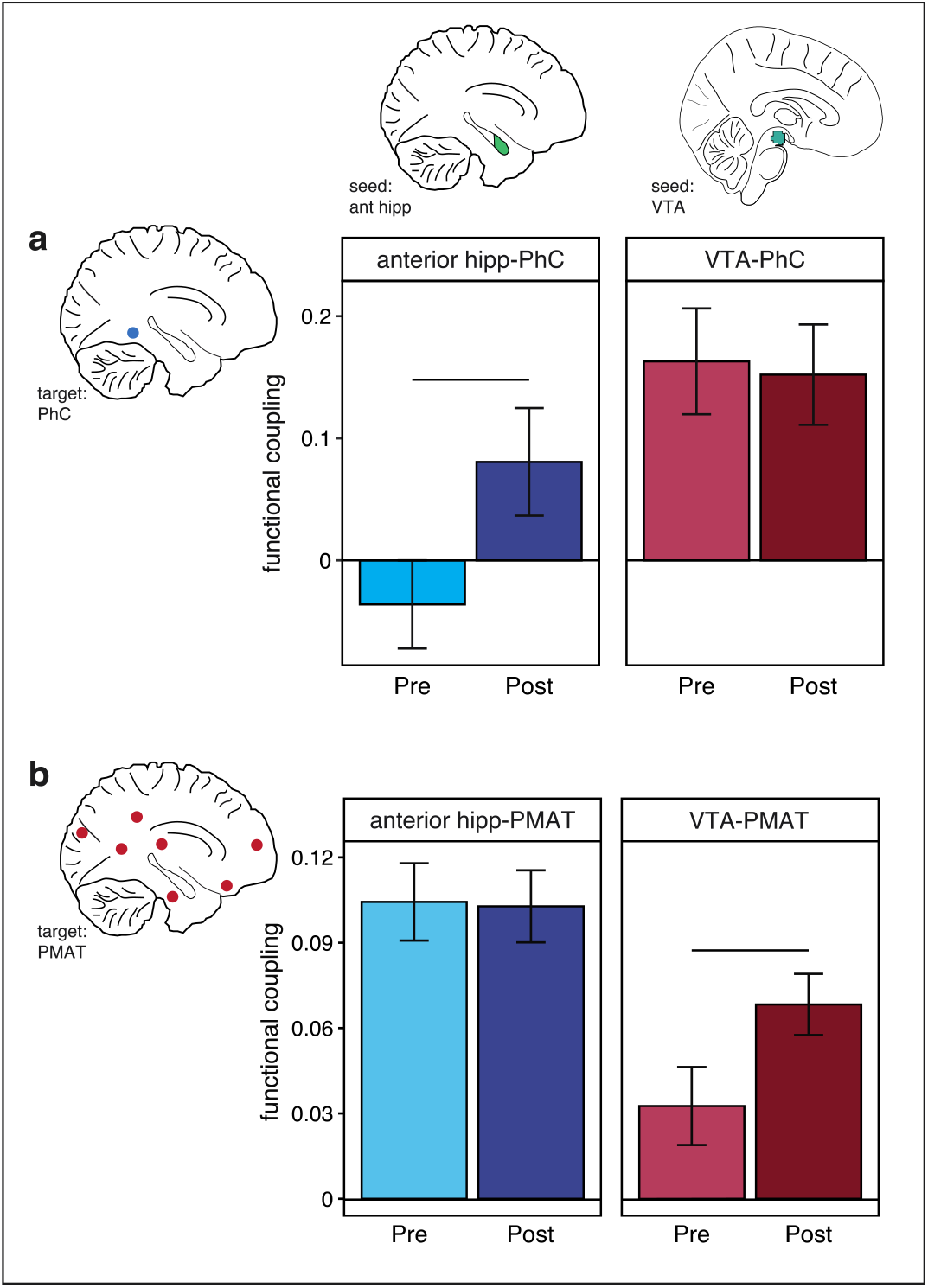
Dissociation in novelty-related plasticity in functional coupling of anterior hippocampus and VTA with cortical network targets. a) Comparing changes in anterior hippocampal (left) and VTA (right) functional coupling with the task-relevant PhC region following novelty-exposure yielded a significant Seed × Session interaction effect, driven by a significant post-novelty enhancement only for anterior hippocampal-PhC functional coupling, with no significant change in VTA-PhC functional coupling from pre to post-novelty exposure. b) In contrast, VTA-PMAT functional coupling showed a post-novelty enhancement compared to pre-novelty baseline, but there was no significant change in anterior hippocampal-PMAT functional coupling, driving the significant Seed × Session interaction effect.

Likewise, a 2(Seed) × 2(Session) repeated measures ANOVA on the functional coupling between anterior hippocampus and VTA with the PMAT network resulted in a significant Seed × Session (*F*(1,36)= 4.63, p=0.038) interaction effect, as well as significant main effects of Seed (F(1,36)=12.63, p= 0.001) and Session (*F*(1,36)=4.58, p= 0.04). However, in contrast to the results with PhC, follow-up t-tests demonstrated this interaction was driven by a significant post-encoding change in functional coupling between VTA and PMAT regions (*t*(36)= −3.81, p=0.0005), with no significant difference in anterior hippocampal-PMAT functional coupling (*t*(36)= 0.11, p=0.91, Figure 3b).

Since regions of the AT network have been implicated in processing affective, salient information (Ranganath and Ritchey, 2012; Ritchey et al., 2015), we separately examined the regions included in the PM and AT systems. As seen in Figure 4, VTA-AT functional coupling showed a significant post-novelty enhancement (*t*(36)= −2.79, p= 0.008), while the change in VTA-PM functional coupling was not statistically significant (*t*(36)= −1.62, p= 0.11). However, as a 2(PMAT network) × 2(Session) repeated measures ANOVA did not yield a significant PMAT network × Session interaction (*F*(1,36)= 0.55, p= 0.46), we continued to collapse across the PMAT networks for further analyses.

**Figure 4.**
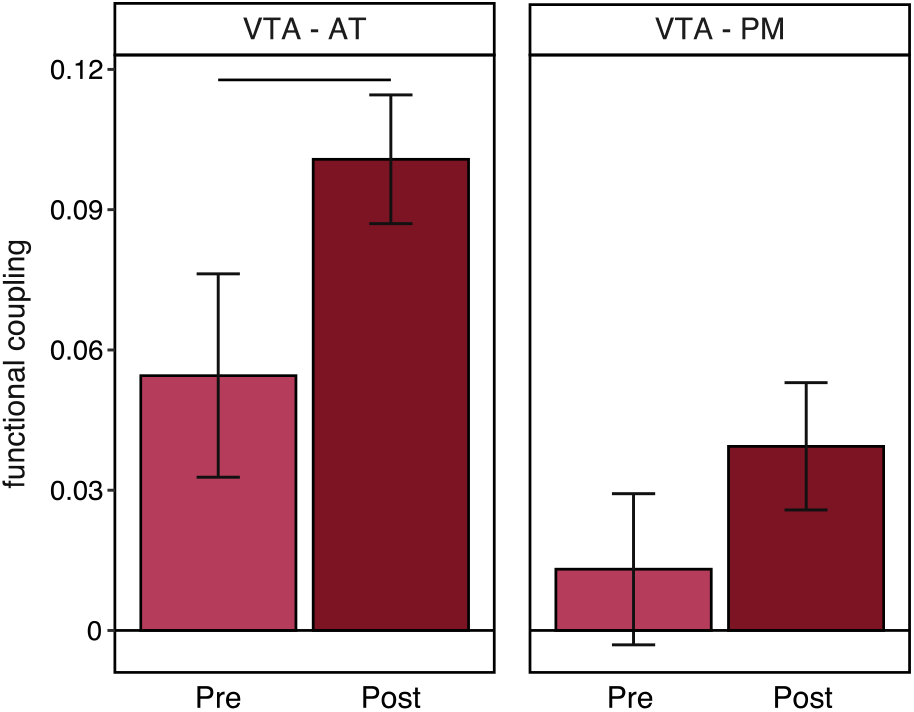
Change in functional coupling between VTA and AT and PM cortical networks. VTA showed significant post-novelty enhancements in functional coupling with the regions of the AT network (left). While not statistically significant, VTA-PM functional coupling showed a similar pattern (right), suggesting the VTA targets relatively diffuse sets of memory-related regions for plasticity-related changes.

To further visualize how widespread the VTA’s post-novelty functional coupling enhancements are with all regions in the PMAT network, we plotted the difference in post – pre-novelty coupling measures between VTA and every region in the PM and AT networks. As Figure 5 illustrates, the majority of regions show positive values, indicating greater post-novelty functional coupling with VTA compared to the pre-novelty measure. In contrast, the same visualization showing the change in functional coupling between anterior hippocampus with all regions of the PMAT networks show more variable patterns (Figure 5). This visualization seems to confirm the consistency with which VTA diffusely targets cortical regions in these networks for novelty-related plasticity benefits.

**Figure 5.**
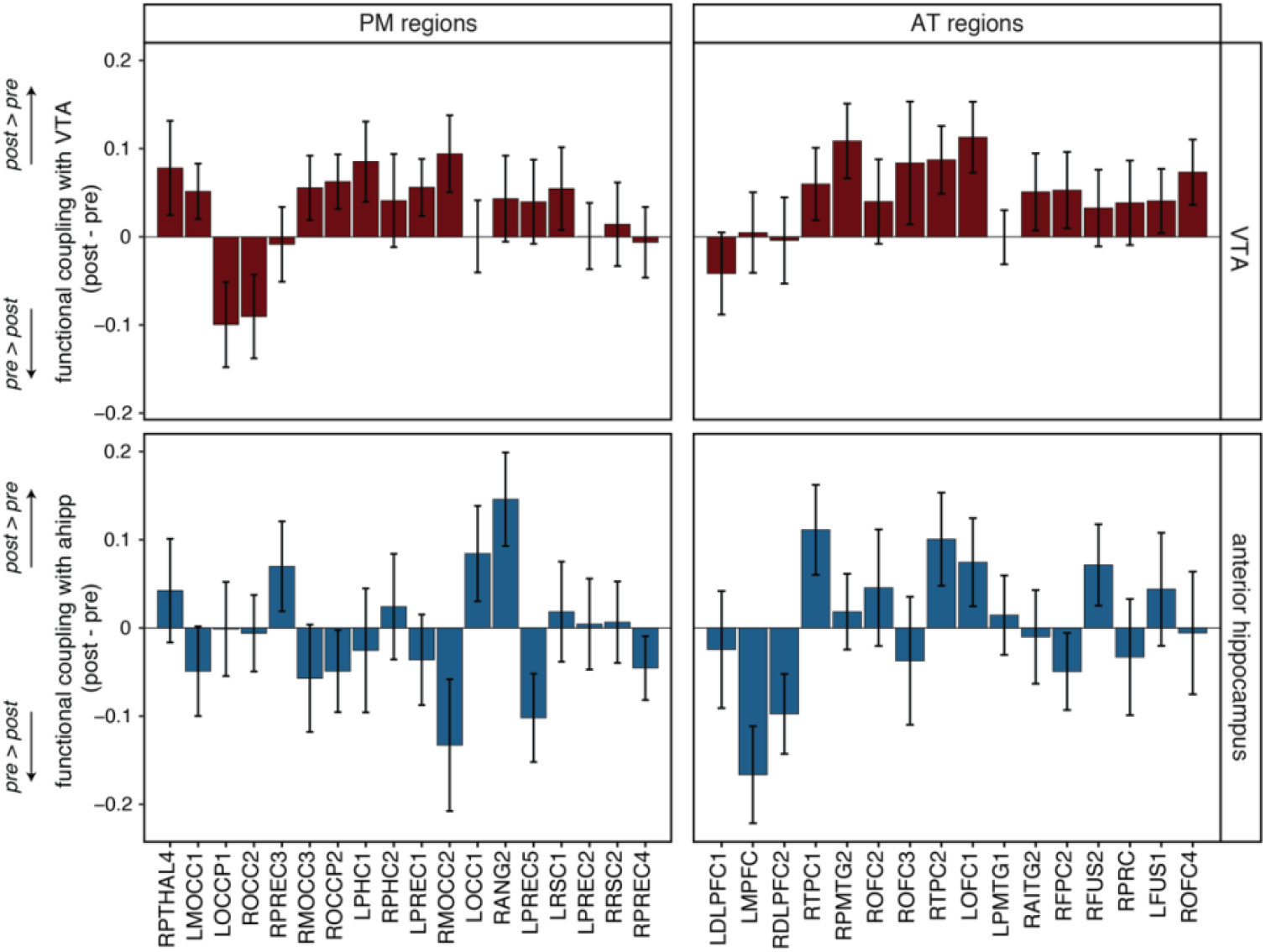
Pairwise change in functional coupling between PMAT regions with VTA and anterior hippocampus. Top: Visualizing the change in functional coupling following novelty exposure (post-pre-novelty) between VTA and all regions included in the PMAT networks illustrates the widespread nature of these effects. The majority of regions show positive difference scores, indicating greater post-novelty functional coupling with VTA compared to the pre-novelty measure. Bottom: In contrast, in the bottom two graphs, the visualization of the post-pre novelty change in functional coupling between anterior hippocampus with all regions included in the PMAT networks shows a variable pattern of effects. Unlike with VTA, only a few regions show positive values, confirming that anterior hippocampus does not seem to target widespread regions for post-novelty functional coupling enhancements. These graphs were used only for visualization purposes, and no statistical tests were calculated on these values.

Together, these results reveal a double dissociation in the cortical networks that anterior hippocampus and VTA target for plasticity-related changes following exposure to novelty, with significant post-encoding enhancements only between anterior hippocampus and the task-relevant parahippocampal cortex and VTA and the diffuse regions in the PMAT network.

### Modulation of plasticity-related changes

To build on these results, we interrogated if this dissociable pattern of novelty-related enhancements in cortical target networks are modulated by the change in the hippocampal-VTA circuit. Since this circuit is central to the detection of novelty and release of dopamine that facilitates plasticity-related changes, we expected that the change in functional coupling between anterior hippocampus and VTA may modulate the extent of plasticity-related changes of each regions’ interactions with the cortex.

We calculated change scores for the functional coupling measures (post-novelty – pre-novelty exposure) and examined the correlations between the change in right anterior hippocampal-VTA functional coupling with the respective changes in right anterior hippocampal and VTA coupling with the diffuse and specific cortical target regions. As shown in Figure 6a, the change in anterior hippocampal-VTA functional coupling positively correlated with both the change in anterior hippocampal-PhC functional coupling (*r*=0.44, p=0.006) and VTA-PMAT functional coupling (*r*= 0.34, p=0.04). Examining this relationship with the networks that did not show significant post-novelty enhancements (Figure 6b), revealed that post-novelty anterior hippocampal-VTA coupling enhancement did not significantly correlate with either the change in VTA-PhC functional coupling (r=0.08, p=0.65) or anterior hippocampal-PMAT coupling (*r*= 0.08, p= 0.66). Moreover, the correlation between the post-novelty enhancements in anterior hippocampal-PhC and VTA-PMAT functional coupling was not statistically significant (*r*=0.16, p= 0.35), potentially indicating a particular role of anterior hippocampal-VTA dynamics in the pathways of cortical network plasticity. Together, these results suggest that the plasticity-related changes resulting from novelty exposure in the anterior hippocampal-VTA network may modulate the effects between anterior hippocampus and VTA with their respective cortical network target regions.

**Figure 6.**
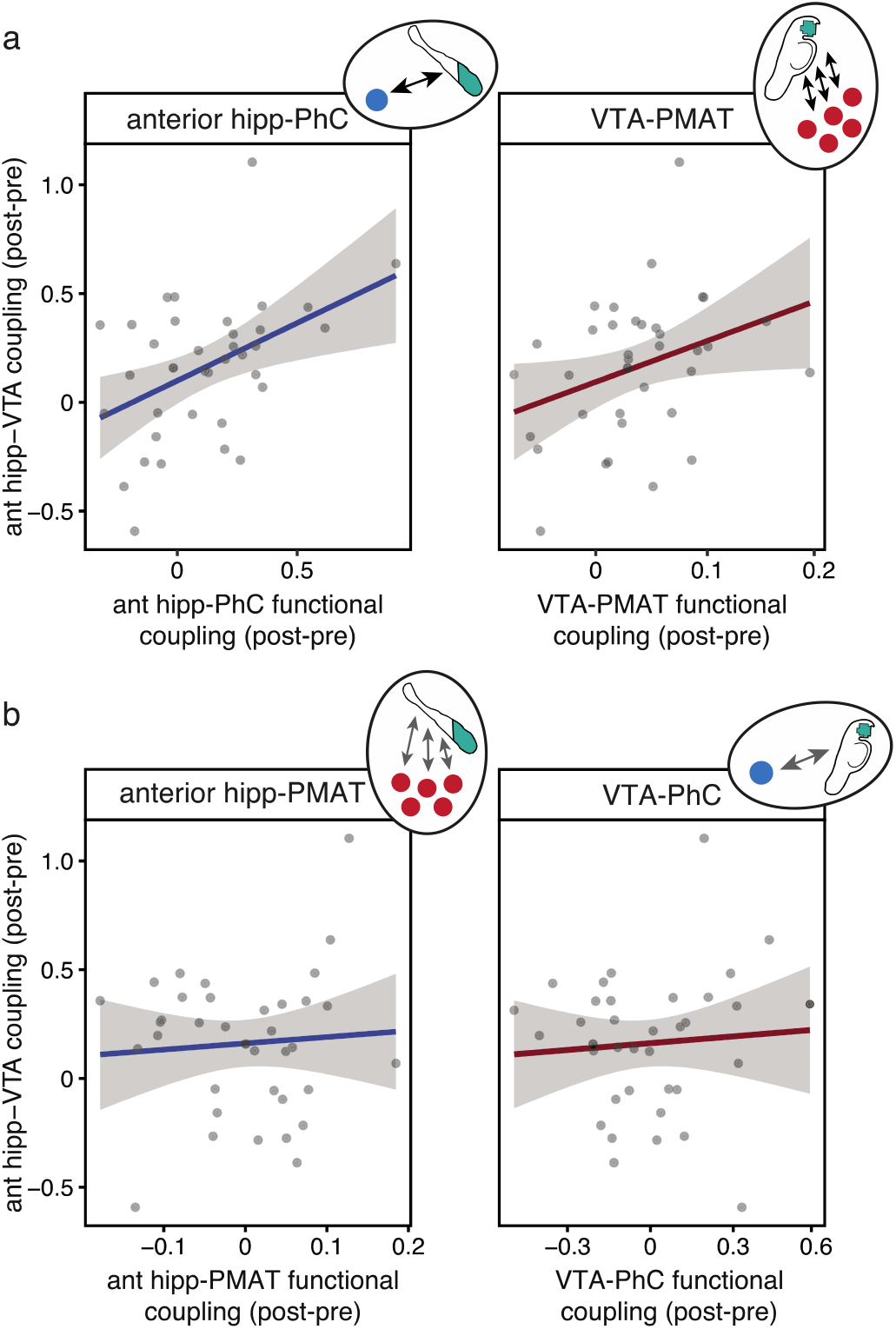
Relationship between anterior hippocampal-VTA functional coupling and coupling with cortical networks. a) The change in anterior hippocampal-VTA functional coupling (post – pre-novelty) positively correlates with the changes in functional coupling between both anterior hippocampal-PhC (left) and VTA-PMAT networks (left), suggesting the extent of novelty-related enhancements in the central anterior hippocampal-VTA circuit is related to the dynamics between anterior hippocampus and VTA with their respective cortical target networks. b) In contrast, the change in anterior hippocampal-VTA functional coupling did not significantly correlate with the change in VTA-PhC (left) functional coupling, nor anterior hippocampal-PMAT functional coupling (right). These results may be in line with the observed scale of distribution in cortical network targets for anterior hippocampus and VTA.

## Discussion

Together, these results provide new evidence that novelty engages cross-regional interactions at differential spatial scales. Comparing changes in resting state functional coupling measured before versus after novelty exposure yielded a series of findings that expand current understanding of the effects of novelty on network-level experience-dependent plasticity. We report several key findings; first, anterior hippocampal-VTA functional coupling was enhanced following novelty exposure, and this change was related to univariate activation in anterior hippocampus during actual exposure to novelty. Further, we found a double dissociation in the cortical regions targeted for plasticity-related changes, with right anterior hippocampus showing experience-dependent increases in functional coupling with a task-relevant region of parahippocampal cortex, but not with the regions of the large-scale PMAT network, while VTA showed enhancements in post-novelty functional coupling with the large-scale memory network, and not task-relevant sensory cortex. These changes in functional coupling correlated with the magnitude of change in anterior hippocampal-VTA functional coupling, suggesting that this central circuit may modulate the plasticity-related changes with cortical regions. These findings provide support for a new model of novelty-induced plasticity, in which novelty elicits plasticity-related changes in both local and global cortical network dynamics.

The hippocampus is sensitive to the detection of novelty (Knight, 1996; Tulving et al., 1996; Strange et al., 1999; Ranganath and Rainer, 2003; Axmacher et al., 2010; Shohamy and Adcock, 2010; Kafkas and Montaldi, 2018), and the underlying hippocampal circuitry may be well suited to facilitate the comparison of incoming sensory inputs with predictions derived from the reactivation of existing prior experiences (Hasselmo and Schnell, 1994; Lisman and Grace, 2005; Kumaran and Maguire, 2006, 2007a, 2007b; Duncan et al., 2012). Potentially due to the unequal proportion of such hippocampal subfields along the long-axis, the anterior hippocampus seems to be functionally specialized for novelty detection (Strange et al., 1999; Poppenk et al., 2013; Kafkas and Montaldi, 2018), consistent with the findings reported here. Once novelty is detected, the hippocampus signals to the VTA to release dopamine, which projects to the hippocampus where it can enhance LTP (Lisman and Grace, 2005). As a result, the synapses coding for the novel information can be strengthened. Accumulating evidence has provided support for this model, with rodent research demonstrating the enhancing effects of dopamine on hippocampal LTP (Huang and Kandel, 1995; Li et al., 2003; Lisman and Grace, 2005; Moncada and Viola, 2007; Rossato et al., 2009; Bethus et al., 2010; Shohamy and Adcock, 2010), and human neuroimaging studies showing increased hippocampal and VTA co-activation and functional coupling during novelty exposure (Schott et al., 2004; Kafkas and Montaldi, 2015; Murty et al., 2017a). We extend evidence for the role of the hippocampus in these processes, demonstrating that functional coupling between the anterior hippocampus and VTA is enhanced following exposure to novelty, indicative of an increase in shared information processing between these regions. Further, this measured change in coupling was related to the univariate activation in the anterior hippocampus evoked during novel trials, in line with the hippocampal detection response facilitating the signaling between these regions.

How does post-encoding facilitation of the VTA-hippocampal circuit propagate changes in cortical network plasticity? Dopamine has been shown to influence replay in the hippocampus during reward motivation (McNamara et al., 2014), suggesting that projections between VTA and hippocampus may also bias the plasticity-related changes instantiated in the cortex. Theories suggest the cortex slowly builds up abstracted representations of information from the hippocampus, resulting in more stable traces that can be integrated into existing stores without interference (McClelland et al., 1995). We interrogated if novelty results in plasticity-related changes in specific task-relevant cortical regions or more diffusely across large-scale cortical networks. Our results suggest that rather than a unified role of the VTA-hippocampal circuit in mediating cortical network plasticity, each region influences the cortex at different resolutions. Specifically, the anterior hippocampus targets local, task-relevant cortical regions, while VTA has more global effects, targeting large-scale networks. Engaging the VTA-hippocampal circuit through novelty exposure may modulate these plasticity-related changes, as we found a positive correlation between the post-novelty change in VTA-anterior hippocampal functional coupling and the change in these respective networks. Together, these data suggest that the effects of novelty extend beyond the hippocampal-VTA circuit, and critically demonstrates that interactions with the cortex are not limited to the hippocampus.

The current findings raise the possibility that there may also be differential consequences of engaging the anterior hippocampal and VTA pathways. Prior reports have shown that similar experience-dependent changes in functional coupling with task-relevant cortical regions are related to enhancements in subsequent memory for the task content (Tambini et al., 2010; Vilberg and Davachi, 2013; Tompary et al., 2015; Murty et al., 2017b; Collins and Dickerson, 2019). It is therefore possible that the hippocampus targets task-relevant regions to stabilize sensory aspects of the novel information to facilitate later memory retrieval. By distributing information processing between the hippocampus and task-relevant regions, novelty could promote the retention of event-specific information.

In contrast, research has implicated the VTA in more diffuse outcomes, with dopaminergic projections from the VTA modulating synaptic plasticity (Jay, 2003; Seamans and Yang, 2004), and influencing the stability of information processing across large-scale cortical networks (Shafiei et al., 2019). Therefore, the post-novelty enhancements in VTA-PMAT functional coupling may reflect more general effects on behavioral processes. Indeed, dopamine seems to play a role in a host of behavioral functions, including memory, decision making, and motor functions (Jay, 2003; Shohamy and Adcock, 2010). Interestingly, there was some indication of a stronger post-novelty enhancement between VTA and the anterior-temporal (AT) network, which has been implicated in mnemonic functions including processing affective and object information (Ranganath and Ritchey, 2012; Ritchey et al., 2015). This is perhaps in line with novelty signaling a motivationally-relevant change in the environment, and facilitating plasticity-related changes to update existing internal models and coordinate behavioral responses. While the current experiment did not include an index of behavioral outcome, future work should explore the behavioral contributions of the interactions between hippocampus and VTA with respective scale of cortical regions.

Novelty is known to have a broad impact on learning, memory, and cognition, by not only facilitating plasticity for novel events, but also by supporting higher-order knowledge that can support exploration, heuristic formation, and schema generation (Krebs et al., 2009; van Kesteren et al., 2012). Yet, it remained unclear how a single behavioral state could facilitate plasticity in a way to foster memory for event-specific details, while also contributing to such higher-order cognition. Our findings begin to illustrate a more complex picture of how novelty influences neural networks to support this diversity in behaviors. We can therefore integrate our findings and prior literature into an extended model, as illustrated in Figure 7. As in prior literature and the Lisman and Grace (2005) model, in this model the hippocampus and VTA form a central, bi-directional circuit, that can detect novelty in the environment, and stimulate dopaminergic release. In addition to effects on hippocampal LTP, activation of this circuit seems to restructure local and global interactions with cortex. The differential scale of distribution in cortical regions targeted by the anterior hippocampus and VTA, with specific task-relevant or large-scale memory networks respectively, suggest that exposure to novelty engages plasticity-related changes across the brain. Such experience-dependent enhancements in interactions between hippocampus and task-relevant sensory cortex may facilitate specific memory reactivation, while VTA coupling with large-scale memory networks may impact information processing across widespread regions. As a result, novel information would be used to update existing models of the world, adaptively facilitating future behavior. Together, this work sheds light on the diverse effects of novelty on functional networks, broadening our understanding of how novelty engages, and restructures, network dynamics across the brain and opening critical questions for future research.

**Figure 7.**
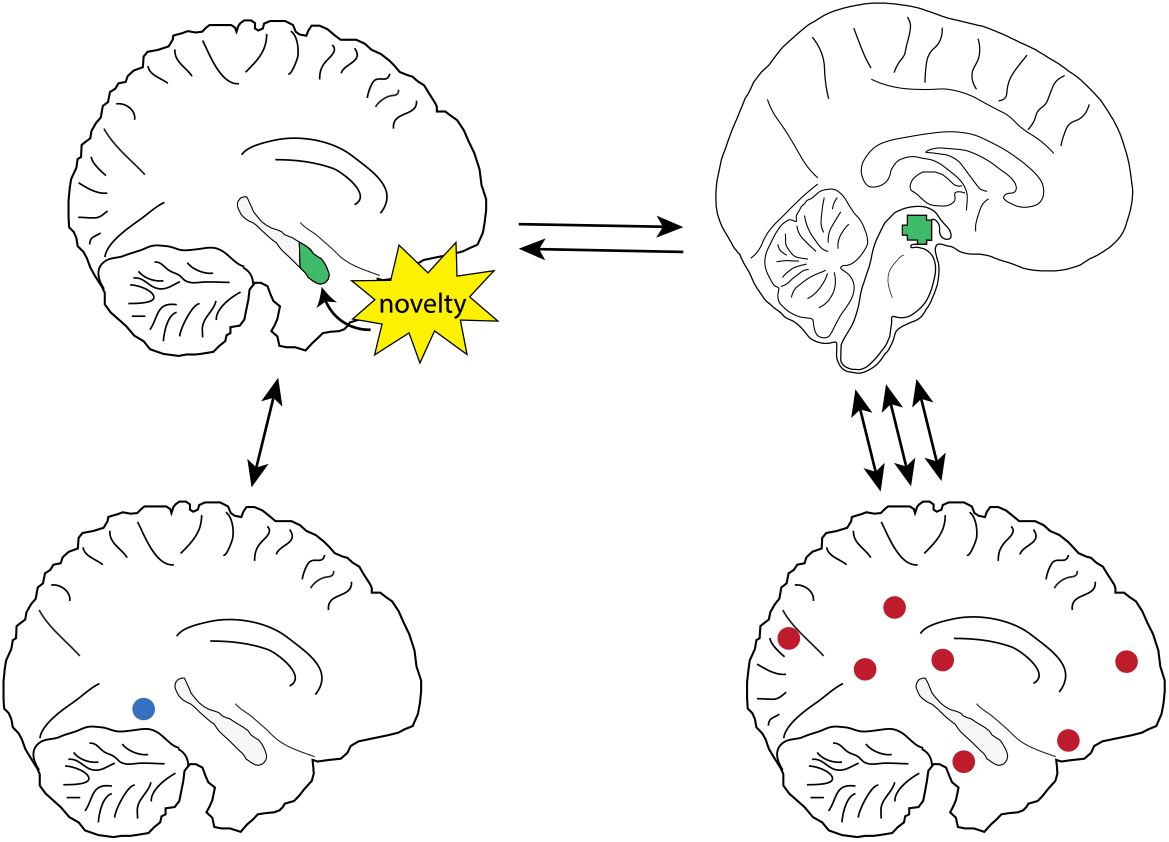
Schematic model illustrating networks engaged through exposure to novelty. In this model integrating the current results with prior literature, the hippocampus and VTA form a bi-directional circuit (top) in which the hippocampus detects novelty in the environment and activates the VTA to release dopamine, in turn enhancing hippocampal LTP. Engaging the VTA-hippocampal circuit also facilitates plasticity-dependent changes in cortical networks, with each region influencing the cortex at distinct scales of distribution. Anterior hippocampus targets task-relevant sensory regions (bottom left) while VTA targets large-scale memory networks (bottom right), potentially contributing to diverse behavioral outcomes evoked by novelty exposure, such as specific memory reactivation and information processing across diffuse regions, respectively. Together this presents an expanded model of how novelty influences neural networks, shedding light on the differential contributions of the anterior hippocampus and VTA to propagating cortical network plasticity.

## Acknowledgements

The project described was supported by a NARSAD Young Investigator Award by the Brain & Behavior Research Foundation awarded to VPM, and the National Institutes of Health through grants K01MH111991 (VPM) and R01MH112613 (LME). We thank Ian Ballard, Alexa Tompary, and James Antony for their comments and feedback on earlier versions of the manuscript.

## References

Axmacher N, Cohen MX, Fell J, Haupt S, Dümpelmann M, Elger CE, Schlaepfer TE, Lenartz D, Sturm V, Ranganath C (2010) Intracranial EEG Correlates of Expectancy and Memory Formation in the Human Hippocampus and Nucleus Accumbens. Neuron 65:541–549.

Barnett AJ, Reilly W, Dimsdale-Zucker H, Mizrak E, Reagh Z, Ranganath C (2020) Organization of cortico-hippocampal networks in the human brain. bioRxiv.

Bethus I, Tse D, Morris RGM (2010) Dopamine and memory: Modulation of the persistence of memory for novel hippocampal NMDA receptor-dependent paired associates. J Neurosci 30:1610–1618.

Collins JA, Dickerson BC (2019) Functional connectivity in category-selective brain networks after encoding predicts subsequent memory. Hippocampus 29:440–450.

Davachi L (2006) Item, context and relational episodic encoding in humans. Curr Opin Neurobiol 16:693–700 Available at: http://linkinghub.elsevier.com/retrieve/pii/S0959438806001528.

Duncan K, Ketz N, Inati SJ, Davachi L (2012) Evidence for area CA1 as a match/mismatch detector: A high-resolution fMRI study of the human hippocampus. Hippocampus 22:389–398.

Girardeau G, Zugaro M (2011) Hippocampal ripples and memory consolidation. Curr Opin Neurobiol 21:452–459 Available at: http://linkinghub.elsevier.com/retrieve/pii/S0959438811000316.

Hasselmo ME, Schnell E (1994) Laminar Selectivity of the Cholinergic Suppression of Synaptic Transmission in Rat Hippocampal Region CA 1: Computational Modeling and Brain Slice Physiology.

Howard LR, Kumaran D, lafsdóttir HF., Spiers HJ (2011) Double dissociation between hippocampal and parahippocampal responses to object-background context and scene novelty. J Neurosci 31:5253–5261.

Huang YY, Kandel ER (1995) D1/D5 receptor agonists induce a protein synthesis-dependent late potentiation in the CA1 region of the hippocampus. Proc Natl Acad Sci U S A 92:2446–2450.

Jay TM (2003) Dopamine: A potential substrate for synaptic plasticity and memory mechanisms. Prog Neurobiol 69:375–390.

Ji D, Wilson MA (2007) Coordinated memory replay in the visual cortex and hippocampus during sleep. Nat Neurosci 10:100–107 Available at: http://www.nature.com/articles/nn1825.

Joo HR, Frank LM (2018) The hippocampal sharp wave–ripple in memory retrieval for immediate use and consolidation. Nat Rev Neurosci 19:744–757.

Kafkas A, Montaldi D (2015) Striatal and midbrain connectivity with the hippocampus selectively boosts memory for contextual novelty. Hippocampus 25:1262–1273.

Kafkas A, Montaldi D (2018) How do memory systems detect and respond to novelty? Neurosci Lett 680:60–68.

Knight RT (1996) Contribution of human hippocampal region to novelty detection. Nature 383:256–259 Available at: http://www.nature.com/articles/383256a0.

Krebs RM, Schott BH, Schütze H, Düzel E (2009) The novelty exploration bonus and its attentional modulation. Neuropsychologia 47:2272–2281.

Kumaran D, Maguire EA (2006) An Unexpected Sequence of Events: Mismatch Detection in the Human Hippocampus Rugg M, ed. PLoS Biol 4:e424 Available at: https://dx.plos.org/10.1371/journal.pbio.0040424.

Kumaran D, Maguire EA (2007a) Match-mismatch processes underlie human hippocampal responses to associative novelty. J Neurosci 27:8517–8524.

Kumaran D, Maguire EA (2007b) Which computational mechanisms operate in the hippocampus during novelty detection? Hippocampus 17:735–748.

Lansink CS, Goltstein PM, Lankelma J V., McNaughton BL, Pennartz CMA (2009) Hippocampus Leads Ventral Striatum in Replay of Place-Reward Information Stevens CF, ed. PLoS Biol 7:e1000173 Available at: https://dx.plos.org/10.1371/journal.pbio.1000173.

Li S, Cullen WK, Anwyl R, Rowan MJ (2003) Dopamine-dependent facilitation of LTP induction in hippocampal CA1 by exposure to spatial novelty. Nat Neurosci 6:526–531.

Lisman JE, Grace AA (2005) The hippocampal-VTA loop: Controlling the entry of information into long-term memory. Neuron 46:703–713.

McClelland JL, McNaughton BL, O’Reilly RC (1995) Why There Are Complementary Learning Systems in the Hippocampus and Neo-cortex: Insights from the Successes and Failures of Connectionists Models of Learning and Memory. Psychol Rev 102:419–457.

McNamara CG, Tejero-Cantero Á, Trouche S, Campo-Urriza N, Dupret D (2014) Dopaminergic neurons promote hippocampal reactivation and spatial memory persistence. Nat Neurosci 17:1658–1660.

Moncada D, Viola H (2007) Induction of long-term memory by exposure to novelty requires protein synthesis: Evidence for a behavioral tagging. J Neurosci 27:7476–7481.

Moscovitch M, Cabeza R, Winocur G, Nadel L (2016) Episodic Memory and Beyond: The Hippocampus and Neocortex in Transformation. Annu Rev Psychol 67:105–134 Available at: http://www.annualreviews.org/doi/10.1146/annurev-psych-113011-143733.

Murty VP, Ballard IC, Adcock RA (2017a) Hippocampus and Prefrontal Cortex Predict Distinct Timescales of Activation in the Human Ventral Tegmental Area. Cereb Cortex 27:1660–1669.

Murty VP, Ballard IC, MacDuffie KE, Krebs RM, Adcock RA (2013) Hippocampal networks habituate as novelty accumulates. Learn Mem 20:229–235.

Murty VP, Dickerson KC (2016) Motivational influences on memory. Adv Motiv Achiev 19:203–227.

Murty VP, Shermohammed M, Smith D V., Carter RMK, Huettel SA, Adcock RA (2014) Resting state networks distinguish human ventral tegmental area from substantia nigra. Neuroimage 100:580–589.

Murty VP, Tompary A, Adcock RA, Davachi L (2017b) Selectivity in Postencoding Connectivity with High-Level Visual Cortex Is Associated with Reward-Motivated Memory. J Neurosci 37:537–545 Available at: http://www.jneurosci.org/lookup/doi/10.1523/JNEUROSCI.4032-15.2016.

Peyrache A, Khamassi M, Benchenane K, Wiener SI, Battaglia FP (2009) Replay of rule-learning related neural patterns in the prefrontal cortex during sleep. Nat Neurosci 12:919–926 Available at: http://dx.doi.org/10.1038/nn.2337.

Poppenk J, Evensmoen HR, Moscovitch M, Nadel L (2013) Long-axis specialization of the human hippocampus. Trends Cogn Sci 17:230–240 Available at: http://www.ncbi.nlm.nih.gov/pubmed/23597720.

Ranganath C, Rainer G (2003) Cognitive neuroscience: Neural mechanisms for detecting and remembering novel events. Nat Rev Neurosci 4:193–202.

Ranganath C, Ritchey M (2012) Two cortical systems for memory-guided behaviour. Nat Rev Neurosci 13:713–726 Available at: http://www.nature.com/articles/nrn3338.

Ritchey M, Libby LA, Ranganath C (2015) Cortico-hippocampal systems involved in memory and cognition. In: Progress in brain research, pp 45–64 Available at: http://linkinghub.elsevier.com/retrieve/pii/S0079612315000588.

Ritchey M, Yonelinas AP, Ranganath C (2014) Functional connectivity relationships predict similarities in task activation and pattern information during associative memory encoding. J Cogn Neurosci 26:1085–1099.

Rossato JI, Bevilaqua LRM, Izquierdo I, Medina JH, Cammarota M (2009) Dopamine controls persistence of long-term memory storage. Science (80-) 325:1017–1020.

Schlichting ML, Preston AR (2014) Memory reactivation during rest supports upcoming learning of related content. Proc Natl Acad Sci 111:15845–15850 Available at: http://www.pnas.org/cgi/doi/10.1073/pnas.1404396111.

Schott BH, Sellner DB, Lauer CJ, Habib R, Frey JU, Guderian S, Heinze HJ, Düzel E (2004) Activation of midbrain structures by associative novelty and the formation of explicit memory in humans. Learn Mem 11:383–387.

Seamans JK, Yang CR (2004) The principal features and mechanisms of dopamine modulation in the prefrontal cortex. Prog Neurobiol 74:1–58.

Shafiei G, Zeighami Y, Clark CA, Coull JT, Nagano-Saito A, Leyton M, Dagher A, Mišić B (2019) Dopamine Signaling Modulates the Stability and Integration of Intrinsic Brain Networks. Cereb Cortex 29:1–13.

Shohamy D, Adcock RA (2010) Dopamine and adaptive memory. Trends Cogn Sci 14:464–472 Available at: https://linkinghub.elsevier.com/retrieve/pii/S1364661310001865.

Squire L, Zola-Morgan S (1991) The medial temporal lobe memory system. Science (80-) 253:1380–1386 Available at: http://www.sciencemag.org/cgi/doi/10.1126/science.1896849.

Strange BA, Fletcher PC, Henson RN, Friston KJ, Dolan RJ (1999) Segregating the functions of human hippocampus. Proc Natl Acad Sci U S A 96:4034–4039 Available at: http://www.ncbi.nlm.nih.gov/pubmed/10097158.

Tambini A, Ketz N, Davachi L (2010) Enhanced Brain Correlations during Rest Are Related to Memory for Recent Experiences. Neuron 65:280–290.

Tompary A, Duncan K, Davachi L (2015) Consolidation of Associative and Item Memory Is Related to Post-Encoding Functional Connectivity between the Ventral Tegmental Area and Different Medial Temporal Lobe Subregions during an Unrelated Task. J Neurosci 35:7326–7331 Available at: http://www.jneurosci.org/cgi/doi/10.1523/JNEUROSCI.4816-14.2015.

Tulving E, Markowitsch HJ, Craik FIM, Habib R, Houle S (1996) Novelty and Familiarity Activations in PET Studies of Memory Encoding and Retrieval. Cereb Cortex 6:71–79 Available at: https://academic.oup.com/cercor/article/6/1/71/265688.

van Kesteren MTR, Ruiter DJ, Fernández G, Henson RN (2012) How schema and novelty augment memory formation. Trends Neurosci 35:211–219 Available at: http://linkinghub.elsevier.com/retrieve/pii/S0166223612000197.

Vilberg KL, Davachi L (2013) Perirhinal-Hippocampal Connectivity during Reactivation Is a Marker for Object-Based Memory Consolidation. Neuron 79:1232–1242 Available at: http://linkinghub.elsevier.com/retrieve/pii/S0896627313006132.

Wang S-H, Morris RGM (2010) Hippocampal-Neocortical Interactions in Memory Formation, Consolidation, and Reconsolidation. Annu Rev Psychol 61:49–79.

Wierzynski CM, Lubenov E V., Gu M, Siapas AG (2009) State-Dependent Spike-Timing Relationships between Hippocampal and Prefrontal Circuits during Sleep. Neuron 61:587–596.

Wilson M, McNaughton B (1994) Reactivation of hippocampal ensemble memories during sleep. Science (80-) 265:676–679 Available at: https://www.sciencemag.org/lookup/doi/10.1126/science.8036517.

